# Enhanced cognitive interference during visuomotor tasks may cause eye-hand dyscoordination

**DOI:** 10.1101/2022.05.03.490541

**Authors:** Tarkeshwar Singh, John R Rizzo, Cédrick Bonnet, Jennifer Semrau, Troy M Herter

**Affiliations:** Department of Kinesiology, The Pennsylvania State University, University Park, PA-16802; Department of Rehabilitation Medicine, New York University Langone Medical Center, New York, NY, USA; Department of Neurology, New York University Langone Medical Center, New York, NY, USA; Univ. Lille, CNRS, UMR 9193-SCALab-Sciences Cognitives et Sciences Affectives, Lille, France; Department of Kinesiology and Applied Physiology, University of Delaware, Newark, DE-19716; Department of Kinesiology, University of South Carolina, Columbia, SC-29208

## Abstract

In complex visuomotor tasks, such as cooking, people make many saccades to continuously search for items before and during reaching movements. These tasks require use of short-term memory and task-switching (e.g., switching search between vegetables and spices). Cognitive load may affect visuomotor performance by increasing the demands on mental processes mediated by the prefrontal cortex, but mechanisms remain unclear. It is also unclear how patients with neurological injuries, e.g., stroke survivors, manage greater cognitive loads during visuomotor tasks. Using the Trail-Making Test, we have previously shown that stroke survivors make many more saccades, which are associated limb movements that are less smooth and slower. In this test, participants search for and make reaching movements towards twenty-five numbers and letters. It has a simple variant (Trails-A), and a cognitively challenging variant (Trails-B) that requires alphanumeric switching. The switching makes the task gradually harder as the Trails-B trial progresses (greater cognitive load). Here, we show that stroke survivors and healthy controls made many more saccades and had longer fixations as the Trails-B trial progressed. In addition, reaching speed slowed down for controls in Trails-B. We propose a mechanism where enhanced cognitive load may reduce inhibition from the prefrontal cortex and disinhibit the ocular motor system into making more saccades. These additional saccades may subsequently slow down motor function by disrupting the visual feedback loops used to control limb movements. These findings augment our understanding of the mechanisms that underpin cognitive interference dynamics when visual, ocular, and limb motor systems interact in visuocognitive motor tasks.

**NEW & Noteworthy:**

o We used a neuropsychological test called the Trails-Making-test and analyze patterns of eye and reaching movements in controls and stroke survivors. We characterized how gaze and reaching movements change within a trial in the easier Trails-A and the more cognitively challenging Trails-B variant that requires alphanumeric switching. We found that as the Trails-B trial progressed participants made more saccadic eye movements and longer fixations, likely because of greater cognitive load.

## Introduction

During activities of daily living, humans continuously re-direct their gaze to search for object(s) and visual stimuli of interest and then initiate actions to interact with them. For example, humans continuously shift gaze while driving amidst traffic lights, street signs, and vehicles before taking appropriate actions at the pedal and steering wheel (1-3). While making a meal, humans continuously search for ingredients and cookware before and during the process of grasping and transporting key items to the right spatial locations for further actions (4). While it is easily taken for granted that our eyes rapidly shift to new locations several times a second (5), visual search is a highly evolved byproduct of evolution and development that is regulated by *top-down* executive processes that interact with bottom-up sensory processes. Top-down executive processes use knowledge from previous experience to direct gaze towards locations in which there is a higher likelihood of finding task-relevant information (e.g., spices would likely be stored together in the same cabinet). *Bottom-up* sensory processes preferentially direct attention and eye movements to salient objects in the visual workspace (e.g., red spice bottle on the white counter). However, the mechanisms that govern the interplay between visual search and limb motor control are not clear.

Numerous studies have examined reaching movements to a single salient target to advance our understanding of the behavioral features of eye-hand coordination (EHC) and their underlying neural mechanisms (6-8). Most research on this paradigm has focused on *bottom-up* processing of sensory information, with a dominant focus on how visual information is gathered and used for motor planning (though cf. (9)). The implicit assumption within this paradigm is that the visual system isolates a single stimulus of interest and performs relevant visuospatial transformations necessary to enable the motor cortex to activate upper limb muscles to bring the limb to the stimulus. However, real-world situations seldom bestow extraordinary saliency to a single stimulus such that it attracts the attention of an actor exclusively towards itself. In contrast, before initialing limb movements in real-world situations, humans must optimally organize their visual search to quickly retrieve task-relevant information from multiple visual stimuli with competing saliency. For example, experienced drivers optimize visual search in busy urban environments (10) and experienced surgeons optimally scan their work environment for task-relevant information compared to inexperienced counterparts (11). For less demanding activities such as meal preparation, humans make saccades to continuously scan for objects of interest before initiating arm and hand movements to manipulate task-relevant objects (2). These studies offer a glimpse into the flexible and dynamic interactions between top-down and bottom-up processes that drive the ocular and the limb motor systems during activities of daily living.

A critical constraint that governs this interaction between the ocular and limb motor systems is that the limbs have much greater inertia than the eyes. As a result, eye movements are far faster and require far less energy than limb movements. This allows humans to make hundreds of saccades (rapid eye movements) per minute, while only committing to a few limb movements when the identities and spatial locations of task-relevant objects have been established with some degree of certainty. In fact, establishing confidence before initiating limb movement is a critical component of skill acquisition in many real-world tasks and occupations, such as driving, flying, surgery, etc. Importantly, the neural mechanisms that govern top-down and bottom-up interactions are likely disrupted in patients with neurological disorders, resulting in suboptimal visual search and deteriorated task performance (12).

We have previously provided evidence supporting this hypothesis using the Trail-Making Test (13), a neuropsychological test used to assess deficits in visual scanning, processing speed, and task-switching. We found that stroke survivors with mild motor impairments executed significantly more saccades than age-matched controls due to deficits in top-down executive processes, such as visuospatial planning and working memory (14). We then showed that these excessive saccades likely interfered with limb movements, leading to worse performance on the Trail-Making Test (15). Specifically, we found that the number of saccades that stroke survivors made during ongoing reaching movements was strongly associated with slower reaching movements and decreased reaching smoothness in stroke survivors. Furthermore, higher number of saccades were correlated with difficulties performing daily tasks involving hand function and mobility (measured using Stroke Impact Scale). This suggests that disruptive interactions between top-down executive processes and the visuomotor system affect functional performance in stroke survivors.

Here, we asked a different question: does an increase in cognitive load that occurs as the trial progresses affect visual search and limb motor control. Alphanumeric switching in Trails-B becomes harder as the trial progresses and that is likely to increase cognitive load. We hypothesized that an increase in cognitive loading (top-down executive processes) would disinhibit the visual system to bottom-up stimulation and cause participants to make more saccades, longer fixations, and slower limb movements. Support for this hypothesis comes from studies that show that when the working memory load is high, the visual system is more sensitive to bottom-up stimulation (16, 17). In our task, this would imply that the gaze will be reflexively directed to more targets in the workspace as the Trails-B trial progresses. Thus, our first prediction was that as the Trails-B trial progresses, the increased cumulative effect of cognitive load would cause more saccades, longer fixations, and slower limb movements, causing a higher degree of eye-hand dyscoordination (EHdC). Our second prediction was that this effect will be stronger in stroke survivors, i.e., stroke survivors will make progressively more saccades and slower limb movements compared to healthy controls. Our results support the first prediction, but not the second prediction. Though, in general, stroke survivors performed worse than controls; eye hand dyscoordination was exacerbated similarly in the two groups as the trial progressed.

## Methods

The experimental setup and the demographics of the participants have been described in detail in our previous publications (14, 15). Briefly, we recruited 16 stroke survivors (mean age 62, range [48,80], 4 females) and 16 control participants (mean age 60, range [52,70], 10 females). Stroke survivors were included if they had suffered a unilateral stroke at least 6 months before testing in either the middle frontal gyrus or the superior parietal gyrus and had difficulty performing one or more relevant activities of daily living (Stroke Impact Scale-16, one or more individual scores <D5). Participants were excluded if they had a history of a central or peripheral neurological disorder (other than stroke) or a musculoskeletal problem of the tested upper extremity. Stroke survivors were also excluded if they exhibited moderate to severe spasticity of the tested upper extremity (Modified Ashworth Scale score ≥2). All participants were screened for visual impairment and visuospatial neglect, and they had no difficulty understanding and following simple instructions. The Institutional Review Board of the University of South Carolina approved the study. All participants provided informed consent before participating in the study.

Experiments were performed on a KINARM Endpoint Lab (KINARM, Canada) integrated with an Eyelink 1000 remote eye tracker (SR Research, Canada) and an augmented reality display. Limb kinematics were sampled at 1000 Hz and low-pass filtered at 20 Hz. Gaze data were sampled at 500 Hz, preprocessed to remove blinks and corneal reflection artefacts, and low-pass filtered at 20 Hz. Saccades and fixations were subsequently identified from the gaze data.

The Trail-Making Test includes two variants, Trails-A in which participants move their hand to draw lines connecting the first 25 positive integers (1, 2, 3, …, 25, Fig. 1A), and Trails-B in which participants draw lines alternating between the first 13 positive integers and the first 12 Roman letters (1, A, 2, B, 3, C, …, 13; Fig. 1B). Trails-A requires participants to search for the next target in the sequence and then reaching to the target once it has been identified. Trails-B is more cognitively challenging because it also requires top-down executive control in the form of working-memory and set-switching to identify the next target in the sequence (13).

**Figure 1:**
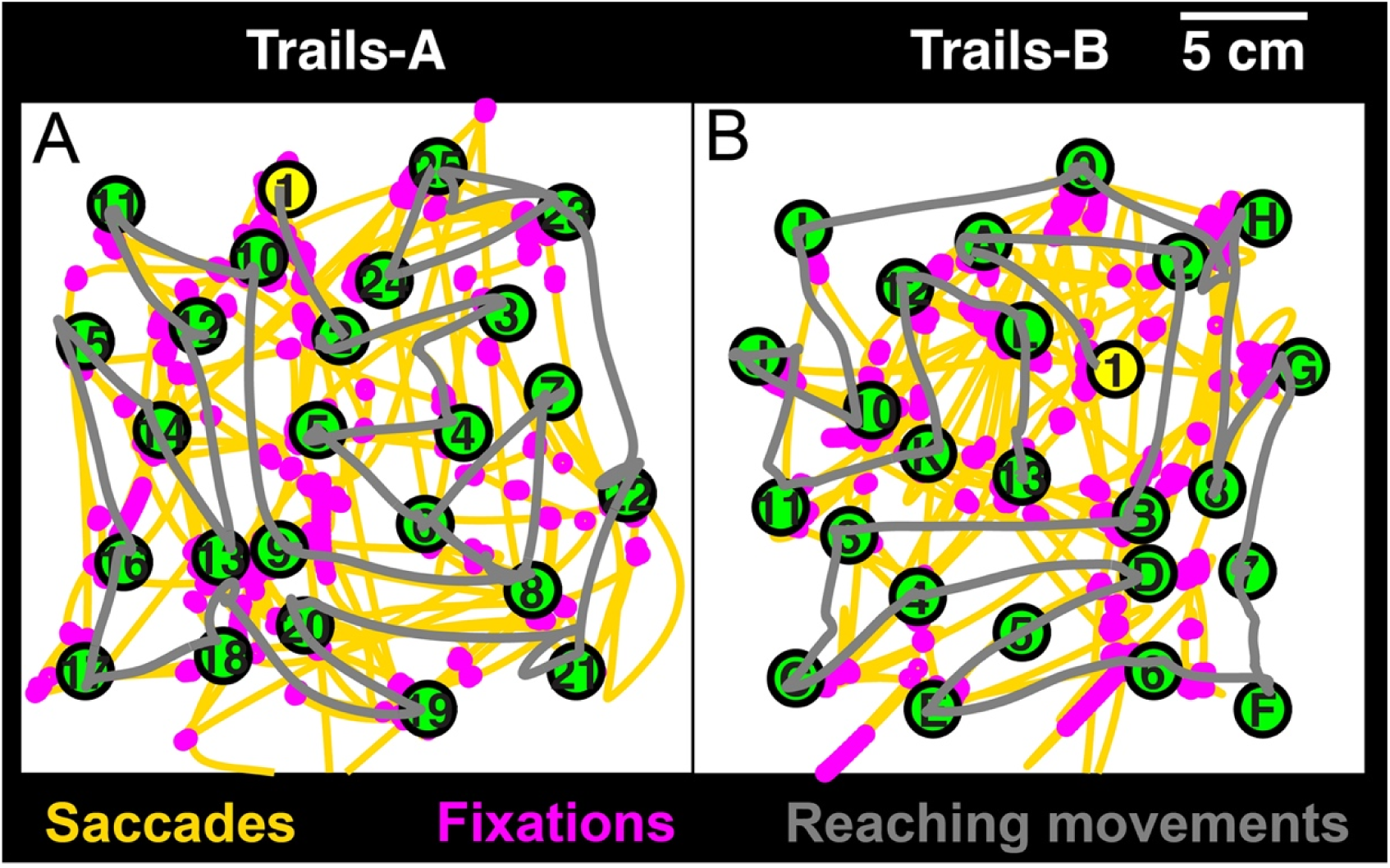
Gaze and hand kinematics during the Trails-Making-Test A (left panel) and B (right panel) for a representative control participant. Saccades are shown in yellow, fixation in pink and the hand reaching movements in grey. Participants cannot see the fixations and saccades, but the reaching trajectories (grey lines) stay on the workspace. Thus, at the very end of the task when only a few targets are left, the grey lines can be used to restrict the visual search in a small area where the remaining targets are.

Participants were instructed to complete the tests as quickly as possible. Importantly, since motor impairments were not the primary interest of this study, stroke survivors were instructed to complete the test with their preferred hand post-stroke and controls with their dominant hand. Most stroke survivors preferred to use their less affected, left hand (post-stroke Edinburgh Handedness Inventory, n = 11). There were no differences in performance between stroke survivors who preferred their left or right hand. Participants performed only a single trial of Trails-A and B within a battery of other eye-hand coordination tasks (not reported here). Before completing the test, participants completed sequences of five practice targets for Trails-A and B.

We examined changes in gaze and reaching behavior as a function of task progression in Trails-A and Trials-B by computing the Number of Saccades, Mean Fixation Duration, and Mean Reach Speed for each sequential pair of targets in Trails-A and Trails-B separately. Measures were quantified from the time the hand touched the first target to the time it touched the second target in each sequential pair. Notably, measures were only quantified for each sequential pair between the 1^st^ and 21^st^ targets. The 21^st^ target was chosen instead of the 25^th^ target due to the spatial layout of the task. Specifically, the search space gradually shrinks between the 1^st^ and 21^st^ targets, but rapidly shrinks between the 22^nd^ and 25^th^ targets. In these small search spaces, both the stroke survivors and controls suddenly produced very few saccades between each sequential pair of targets.

Descriptive statistics are presented in the text and figures as mean ± SE. We performed regressions between the measures and targets for stroke survivors and controls in Trails-A and Trails-B separately. A slope was considered statistically indistinguishable from ‘0’ If its 95% confidence interval spanned both negative and positive numbers. Finally, we used repeated-measures ANOVA with *group* as between-subjects factor (controls and stroke survivors as levels) and *pairs* as the within-subjects factor (first target pair 1-2 and second target pair 20-21 as levels) for Trails-A and Trails-B separately. The level of significance was chosen as a=0.05. Effect sizes were calculated using generalized *η*^2^. Normality of the data were tested using the Wilk-Shapiro test. Bonferroni corrections were used for multiple comparisons. All data pre-processing and analysis were performed in MATLAB (Mathworks, Natick, MA). All statistical tests were performed in R.

## Results

We have previously shown that in both versions of the Trails-test, stroke survivors make more total saccades (and fixations) than age-matched controls (Fig. 5 in 14). Here, we show that for both Trails-A and Trails-B, the number of saccades made by stroke survivors per movement increased as the trial progressed (Fig. 2A). On average, controls made 4.3±1.4 saccades and 3.1±1.0 between the first and last target pairs, respectively, in Trails-A. Stroke survivors made 1.94±0.43 and 6.8±1.29 for the first and the last pairs, respectively. The slope of the regression line was negative for the controls (−0.1, 95% interval [-0.15, -0.05]), and statistically indistinguishable from 0 for stroke survivors. The intercepts were similar for the controls (4.5, 95% interval [3.9,5.1]) and the stroke survivors (4.7, 95% interval [3.0,6.3]). The statistical model showed a significant main effect of *pairs* (p=0.018, *η*^2^=0.08) and an interaction effect between *group* and *pairs* (p=0.007, *η*^2^=0.11). Post-hoc comparisons revealed significant differences between the two groups at the second target pair (p=0.03) and between the first and second targets pairs for the stroke survivors (p=0.002) (Fig. 2B).

**Figure 2:**
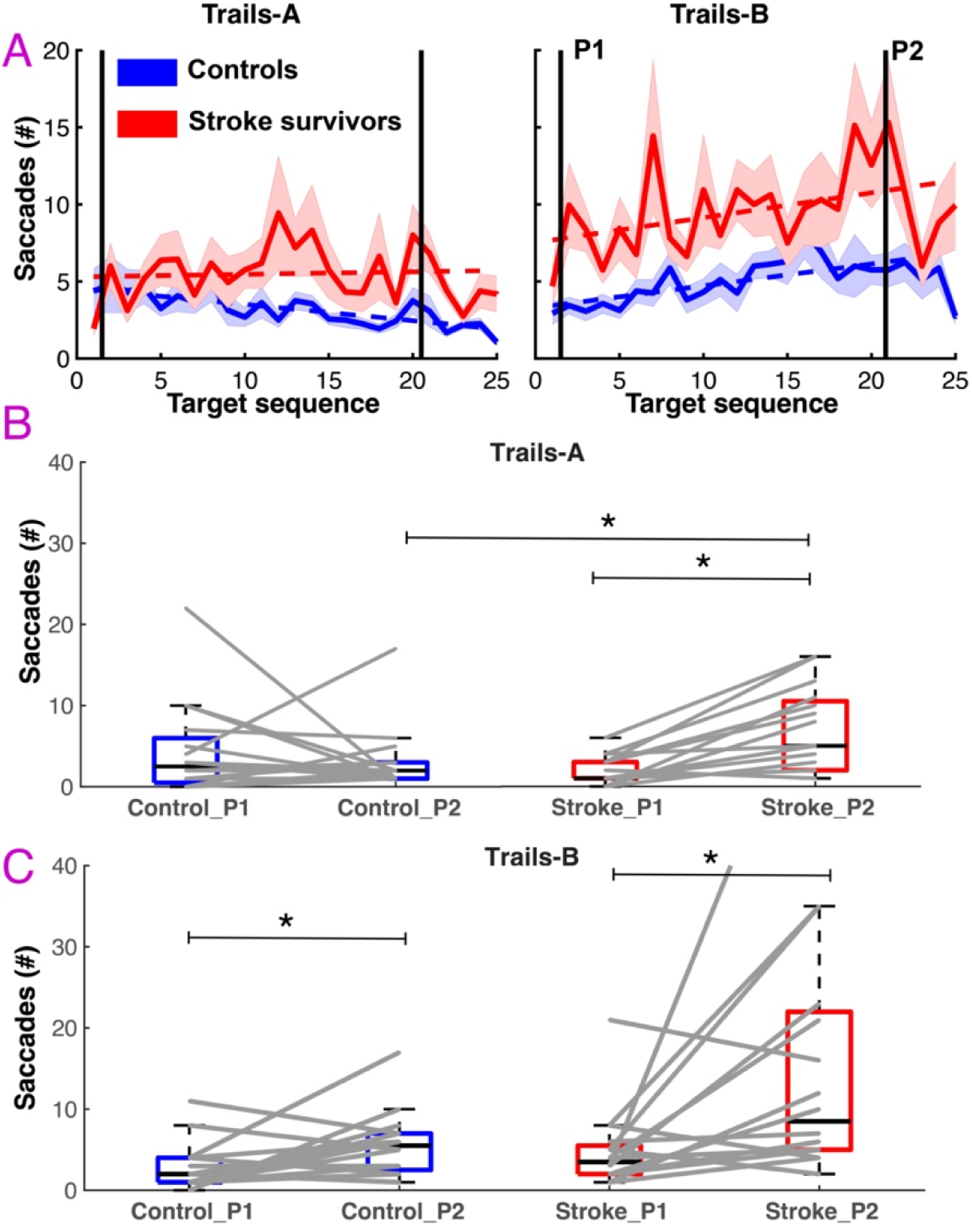
The number of saccadic eye movements increased as the Trails-B trial progressed. A) A regression model fit to the data between the first pair (P1) and last pair (P2) of targets shows no increase in number of saccades as the trial progressed in Trails-A. The number of saccades made per target increased for both controls and stroke survivors in Trails-B, but the regression slope was marginally steeper for stroke survivors. B) Boxplots for Trails-A shows that the stroke survivors made significantly more saccades than controls at the last target pair (P2). Stroke survivors also made more saccades at the last target pair (P2) than the first target pair (P1). C) Boxplot for Trails-B shows that both groups made more saccades at the last target pair (P2) than the first target pair (P1).

In Trails-B, controls made 3.0±0.73 saccades and 4.8±1.2 between the first and last target pairs, respectively. Stroke survivors made a slightly higher number of saccades for the first pair, 4.8±1.2, but many more saccades for the final pair, 16.1±4.3. The intercept was larger for the stroke survivors, and the slope was marginally steeper. The intercept for controls was 3.0 (95% interval [2.1,3.8]) and for the stroke survivors 6.6 (95% interval [4.4,8.8]). The slope for the controls was 0.18 (95% interval [0.11,0.25]) and for the stroke survivors was 0.31 (95% interval [0.13,0.49]) (Fig. 2A). The main effects of *group* (p=0.011, *η*^2^=0.12) and *pairs* (p<0.001, *η*^2^=0.23) were significant. Post-hoc tests showed significant differences between the target pairs for controls (p=0.048) and stroke survivors (p=0.011) (Fig. 2C). Together, these results suggest that both groups made more saccades as the trial progressed in Trails-B, but only stroke survivors made more saccades as the Trails-A trial progressed.

We found very similar results for Mean Fixation Duration (Fig. 3). For controls, the mean fixation duration for the first pair in Trails-A was 0.63±0.25 s and for the last pair was 0.24±0.29 s. For stroke survivors, the fixation duration for the first and final pairs were 0.31±0.11 s and 2.19±0.41 s, respectively. The intercept for the regression was slightly lower for the controls (1.03, 95% interval [0.83,1.25]) than the stroke survivors (1.53, 95% interval [1.0,2.07]). Both the slopes were statistically indistinguishable from 0 (Fig. 3A). The statistical model showed a main effect of *pairs* (p<0.001, *η*^2^=0.28) and an interaction effect between *group* and *pairs* (p=0.014, *η*^2^=0.09). The post-hoc tests showed a significant increase in fixation duration for stroke survivors between the two pairs (p<0.001) (Fig. 3B).

**Figure 3:**
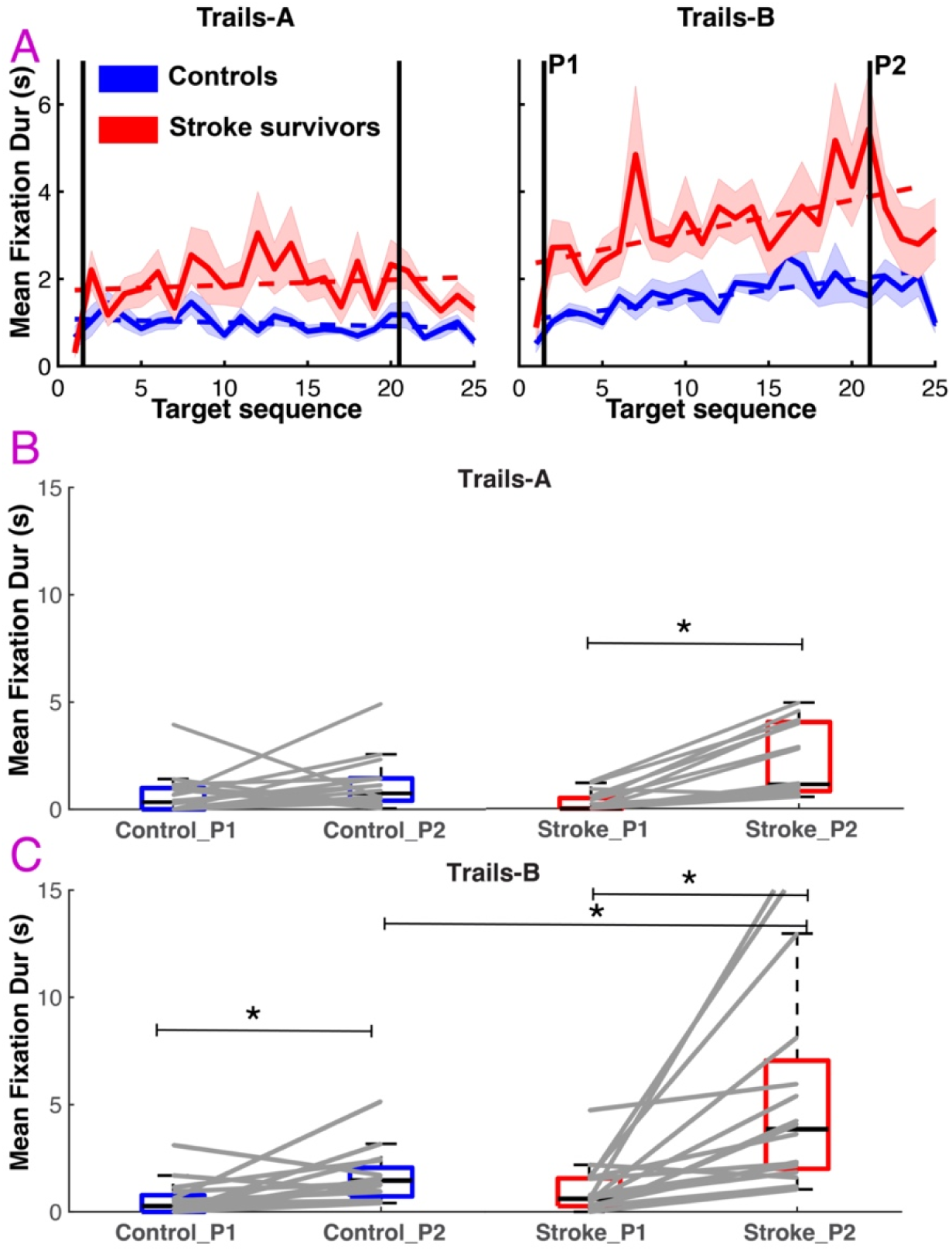
The mean fixation duration increased as the Trails-B trial progressed. A) A regression model fit to the data between the first target pair (P1) and last pair (P2) of targets shows no increase in mean fixation duration as the trial progressed in Trails-A. The mean fixation duration increased for both controls and stroke survivors in Trails-B, and again the regression slope was marginally steeper for stroke survivors suggesting they made longer fixations as the Trails-B trial progressed. B) Boxplots for Trails-A shows that the stroke survivors made significantly longer fixation than controls at the last target pair (P2). C) Boxplot for Trails-B shows that both groups made longer fixations at the last target pair (P2) than the first target pair (P1). Stroke survivors also made more saccades at the last target pair (P2) than the controls.

For Trails-B, mean fixation duration for the controls were 0.56±0.2 s and 1.62±0.3 s for the first and final pairs, respectively. For stroke survivors, the fixation durations were 1.04±0.3 s and 5.6±1.3 s, respectively. The intercept was higher for the stroke survivors (1.99, 95% interval [1.27,2.71]) than controls (0.98, 95% interval [10.68,1.29]). The slopes for the controls (0.06, 95% interval [0.03,0.08]) and stroke survivors were both positive (0.12, 95% interval [0.06,0.18]), but marginally steeper for stroke survivors. Our statistical model showed that the main effects for *group* (p<0.001, *η*^2^= 0.18) and *pairs* (p<0.001, *η*^2^= 0.41) were significant and so was the interaction effect (p=0.017, *η*^2^= 0.07). Post-hoc tests showed significant differences between the two pairs for controls (p=0.002) and stroke survivors (p<0.001) (Fig. 3C). The fixation duration was also significantly higher for the stroke survivors at the second pair (p=0.005). Together, these results suggest that both groups made longer fixations as the trial progressed in Trails-B, but stroke survivors made longer fixations than controls. Stroke survivors also made longer fixations at the second pair in Trails-A.

The Mean Reach Speed (Fig. 4) for Trails-B appeared to decrease for both groups as the trial progressed, but the results were not unequivocal. In Trails-A, the Mean Reach Speed for controls at the first pair of targets was 14.8±1.25 cm/s and for the last pair was 16.4±1.3 cm/s. For stroke survivors, the Mean Reach Speed was 13.1±1.7 cm/s and 11.5±1.6 cm/s, respectively. However, the regression intercepts were very similar between the two groups and the slopes were statistically indistinguishable from 0 (Fig. 4A). For Trails-A, the main effect of *group* was significant (p=0.025, *η*^2^= 0.07). No other effects were significant and no post-hoc tests were significant (Fig. 4B).

**Figure 4:**
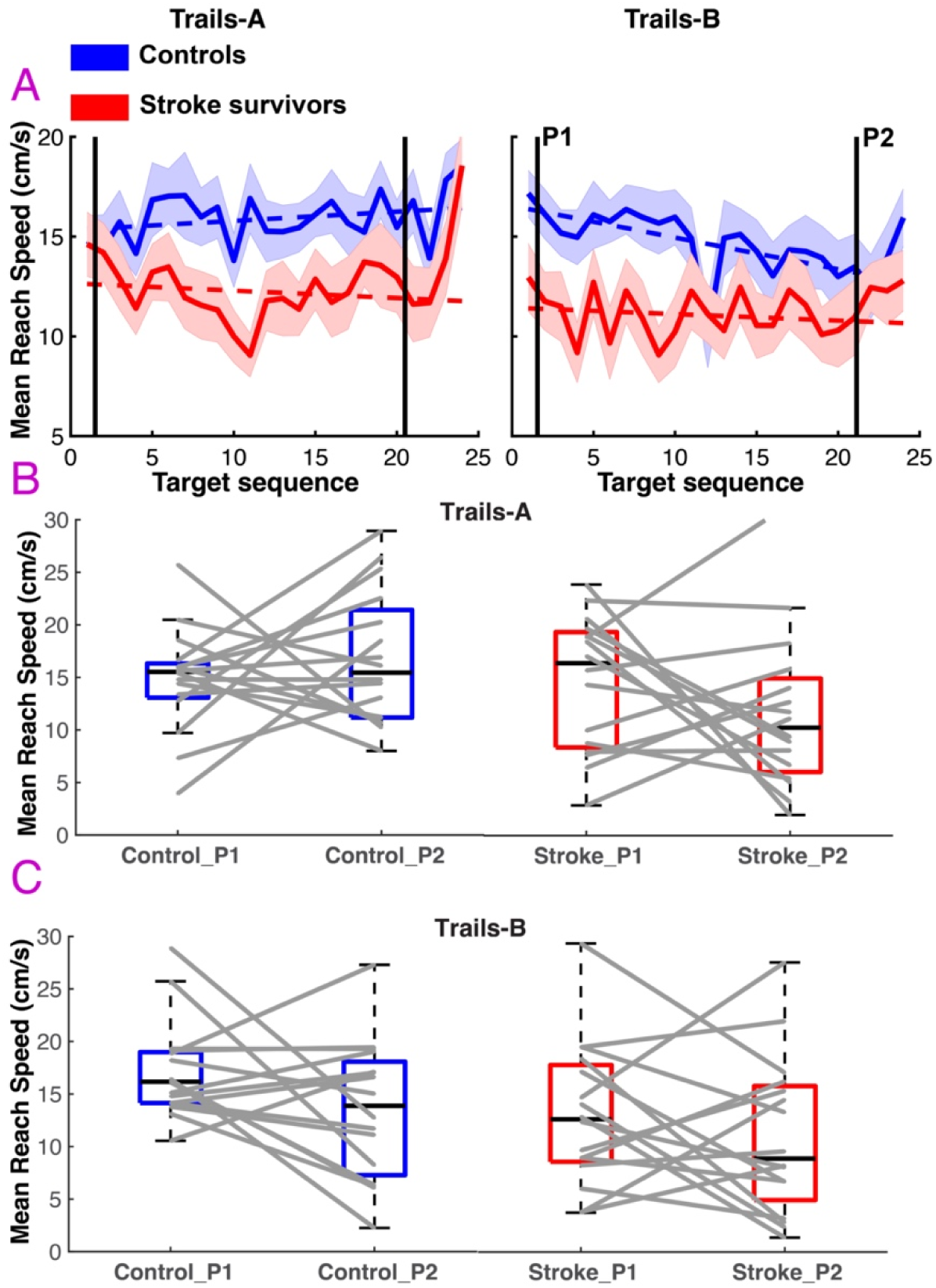
The mean reach speed appears to decrease for controls in Trails-B. A) A regression model fit to the data between the first target pair (P1) and last pair (P2) of targets shows no change in mean reach speed as the trial progressed in Trails-A. The mean reach speed appeared to decrease for both controls and stroke survivors in Trails-B, but the 95% confidence interval of the slope was negative only for the control group. In contrast, the intercepts were significantly lower for the stroke survivors in Trails-B, suggesting that the stroke survivors were overwhelmed with Trails-B throughout the entire task. B) Boxplots for Trails-A shows no significant differences. C) Boxplot for Trails-B shows no significant differences.

For Trails-B, the Mean Reach Speed for the first and final pair of targets for the controls were 15.3±1.3 cm/s and 12.5±2.23 cm/s, respectively. For stroke survivors, the Mean Reach Speed was lower. For the first pair it was 11.3±1.9 cm/s and for the final pair, 7.4±1.98 cm/s. The intercept of the regression was significantly lower for stroke survivors (10.6, 95% interval [9.02,12.12]) than the controls (16.16, 95% interval [14.8,17.4]). The regression slope for controls was -0.16 (95% interval [-0.25, -0.06]). The slope for stroke survivors was also negative (−0.03), but statistically indistinguishable from 0 (Fig. 4A). The statistical model showed a main effect of *group* (p=0.027, *η*^2^= 0.09) and *pairs* (p=0.035, *η*^2^= 0.07). The post-hoc tests revealed no significant differences (Fig. 4C). Together, these results suggest that the Mean Reach Speed was slower for the stroke survivors, but the speed did not change for either group in Trails-A. For Trails-B, the Mean Reach Speed for the stroke survivors was also much slower than the control in Trails-B. The reaching speed decreased more for controls as the trial progressed, but the decrease was not significant.

Finally, we correlated the number of saccades made per movement with the Mean Reach Speed per movement and found that they were not significantly correlated for either the controls or stroke survivors in Trails-A (Fig. 5A). In Trails-B, however, the correlation was significantly negative for the controls (r=-0.52, p<0.05), but not for the stroke survivors (r=-0.24, p=0.2, Fig. 5B). The negative correlation suggests that the higher the number of saccades made for a target pair, the slower was the average reaching speed between those targets. For stroke survivors, the correlation was not significant for Trails-B, and this suggests that other visuomotor factors could have played a role in slowing down of the reaching movements.

**Figure 5:**
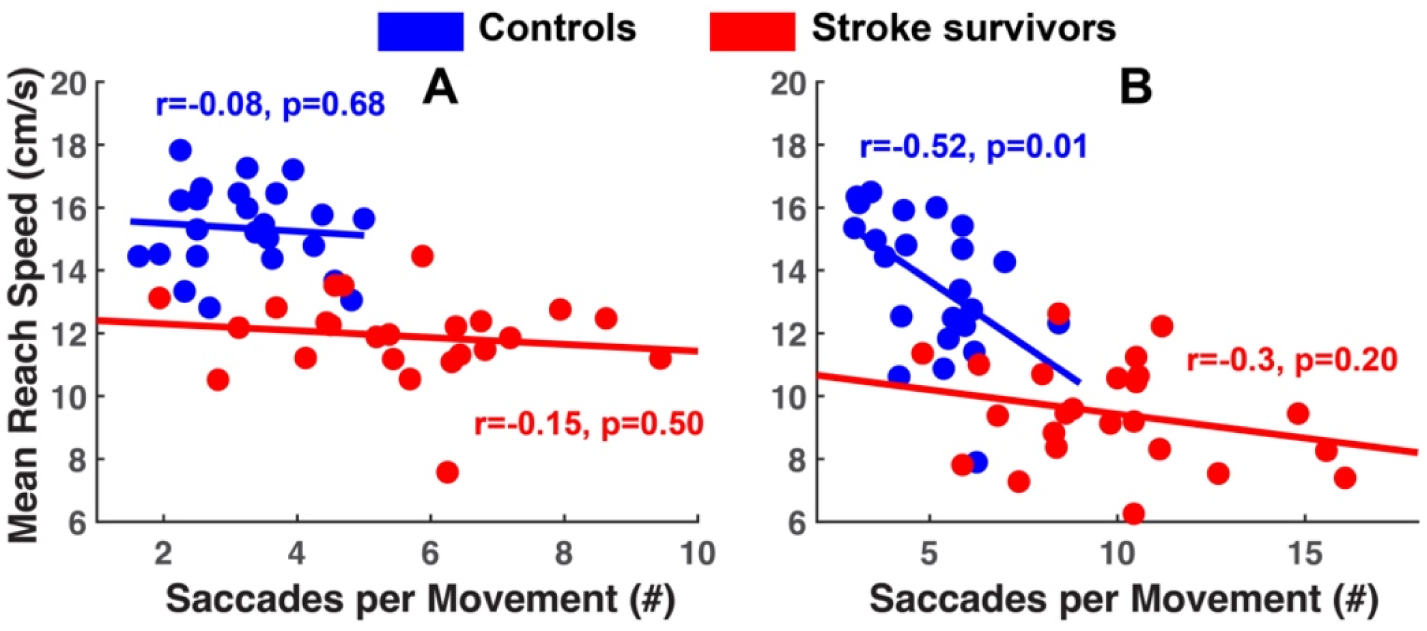
Mean Reach Speed versus number of Saccades made per reach show stronger negative correlations during Trails-B (panel B) than Trails-A (panel A). There is no correlation between these two variables in Trails-A. In Trails-B, the correlation is only significant for controls. However, the stroke survivors made much slower movements despite performing the task with their less impaired hand.

## Discussion

Overall, our results partially support our predictions. Our first prediction was that as the Trails-B trial progressed, the increased cognitive load would cause more saccades, longer fixations, and slower limb movements. This would suggest a higher degree of eye-hand dyscoordination (EHdC). We found that the number of saccades and the mean fixation duration increased as the trial progressed, and that the reaching speed decreased slightly for controls. However, the evidence to support a decrease in reaching speed was not as strong as the evidence for saccades and fixations. Only the regression slope for reaching speed in Trails-B trials was negative for controls, but the statistical model did not confirm this. Our second prediction was that EHdC will be stronger in stroke survivors, i.e., stroke survivors will make progressively more saccades and slower limb movements compared to healthy controls. Stroke survivors made more saccades and longer fixations in Trails-B, but their reaching speed (which was much worse than the controls throughout the Trails-B trial) did not further deteriorate significantly. Previously, we have shown that average number of saccades made per reaching movements were strongly correlated with lower scores on the Stroke Impact Scale (Hand & Mobility section) (15). Together, these results suggest that eye hand dyscoordination (EHdC) may be exacerbated in stroke survivors when they perform visuo-cognitive tasks where the cognitive load increases as a function of time or task difficulty.

In contrast to Trails-A, Trails-B also requires top-down executive processes such as working memory and set-switching. We found that number of saccades and average fixation duration increased, and average reaching speed decreased, as the task progressed in Trails-B. Interestingly, when the task became easier again at the end (limited search space), the number of saccades and fixation durations decreased for stroke survivors with reciprocal increases in hand speed for stroke survivors (see Figs. 2-4). Furthermore, the number of saccades were negatively correlated with reaching movement speed for controls in Trails-B. Though the reaching speed for stroke survivors did not decrease significantly as the Trails-B trial progressed, the reaching speed in general was much slower than controls and relatively slower to earlier reaches even though they performed the test with their preferred and/or less-affected hand. This suggests that stroke survivors were overwhelmed by the Trails-B task, which resulted in poorer task performance.

While an increase in eye movement behavior has been clearly documented in eye-hand tasks in chronic stroke survivors (14, 15, 18, 19), the striking findings in the current study were the large number of saccades that stroke survivors made during reaching movements throughout Trials-B and the increase in number of saccades made by both the controls and stroke survivors during reaching movements as Trails-B progressed. Notably, during Trails-B, stroke survivors started at 4.8 saccades per reach on average and dramatically increased to 16.1 saccades per reach on average compared to an increase from 3.0 to 4.8 saccades per reach on average for the controls. These findings highlight that the difficulties performing rapid movements and preventing unwanted movements are critical but unrecognized aspects of motor control in stroke survivors and, to a lesser extent, controls (20).

Our findings also underscore a potential pathophysiological process that may underlie eye-hand dyscoordination (EHdC) with increasing cognitive demands in stroke survivors and controls. We posit that disinhibition of the ocular motor system during visual search (19) creates downstream interference on limb movements, likely through pathways involving the basal ganglia (21, 22) (see Fig. 6). There is ample evidence to support that the prefrontal cortex is comprised of distinct and overlapping networks involved in response inhibition, task-switching, and working memory (23). Competitive interactions between these networks may result in mutual inhibition, such that increased demands on task switching in Trails-B may interfere with response inhibition and working memory. The effects could include (1) disinhibition of eye movements and (2) difficulty using working memory to guide visual search. Furthermore, these excessive eye movements would overload the visual processing system. This could indirectly inhibit motor function through changes in cortical interactions between the parietofrontal visual networks and the premotor cortex and the subcortical projections from the superior colliculus to the anterior thalamus and motor cortex (22).

**Figure 6:**
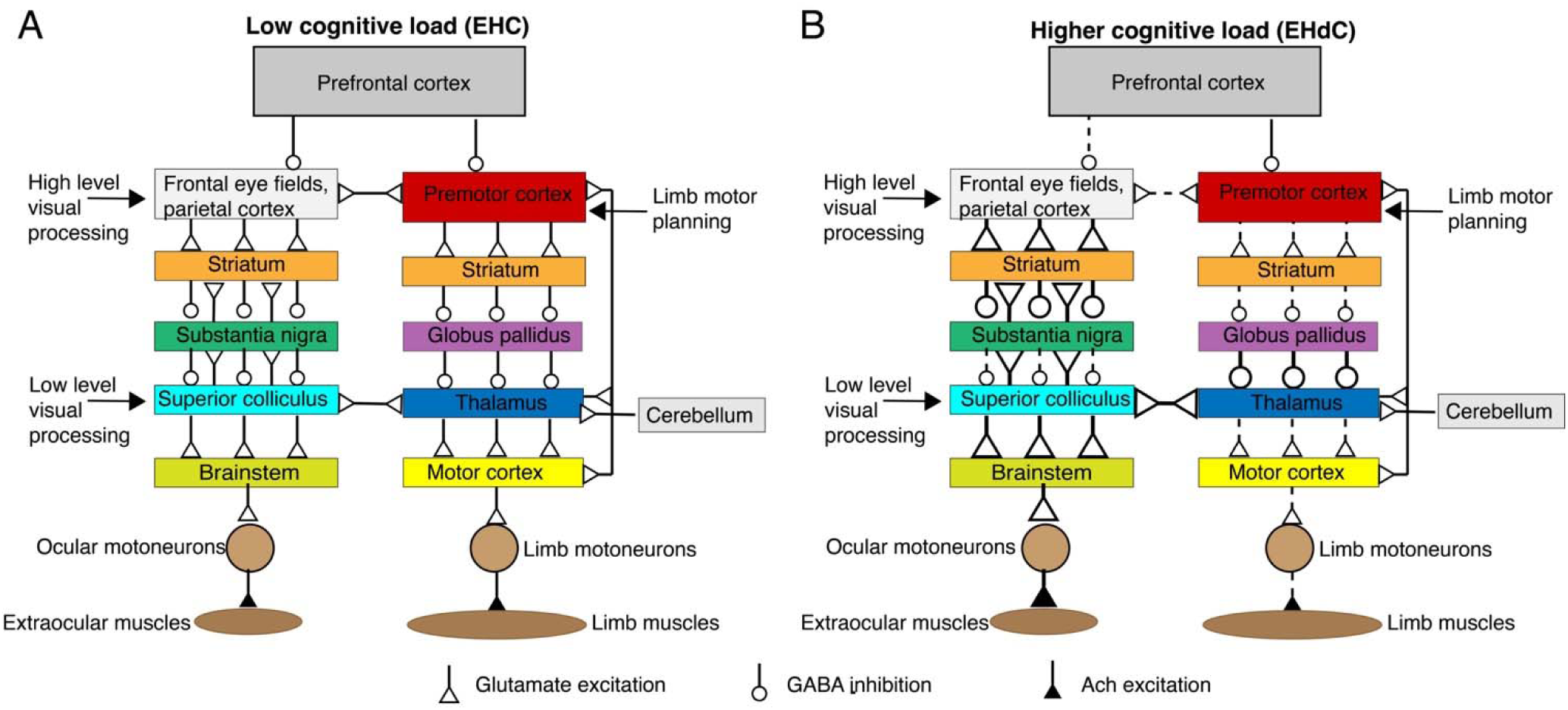
Proposed model for eye hand dyscoordination (EHdC) under enhanced cognitive load. Panel A shows the visual and ocular motor networks involved in eye movements (left) and limb movements (right) along with excitatory and inhibitory projections. The feedback projections within the visual system and interconnections between the visual, ocular motor, and limb motor systems are also shown. We propose that during enhanced cognitive loading, competitive interactions between prefrontal networks involved in response inhibition, task-switching, and working memory disinhibit the parietofrontal areas involved in ocular motor control. This increases the number of saccadic eye movements. The higher number of eye movements likely overload the visual system. This is supported by the longer fixation durations we observed in Trails-B as the task progressed. Then by changes in cortical and subcortical interactions between the ocular motor and limb motor systems, the motor system may also be inhibited during increased cognitive loading. Dashed lines indicate weakened projections and wider lines with larger arrow heads indicate stronger projections.

Fixation durations are another sensitive marker of memory load and processing load, and they are more sensitive to other biomarkers, such as pupil size (24), fixation duration has not been well studied in stroke physiology. Herein, we clearly demonstrate large differences between control and stroke not only in total magnitude of duration, with stroke survivors increasing the total length of a duration up to 4-fold but also in the time course of the increases with fixational duration increasing over the course of the task in Trails-B. In visual search and free-viewing, it is generally well-established that fixational durations increase in length, as the searcher moves from a global to a more focused and local search strategy. This trend is typically observed over the first few seconds of visual search behavior (25). It has also been shown that in multi-target visual search, fixational durations generally increase with increasing memory load (24). In the case of Trail Making Test, a unique global to local strategy is required over a sequential multi-target task until all the numbers and/or letters are acquired. Fixation durational may indicate both the amount of information being processed and also the difficulty of future saccadic target selection (26).

Reaching speed was clearly different between controls and stroke survivors. It should be noted again that most stroke survivors used the ipsilesional limb (i.e., the less affected arm). Although some deficits in the ipsilesional limb have been documented (27, 28), leading to impaired motor function, the striking findings herein was the time course of reaching deficits, which are exacerbated over the course of the trial. In the correlations, for both controls and stroke, interference scales in a stepwise manner for each additional eye movement made. While stroke survivors show a small decrement in reaching performance, controls showed precipitous declines in reaching speeds (see Fig. 4). Previously, it had been demonstrated that stroke survivors made more saccades per reach if more impaired functionally (Fig. 4D in 15) and that these saccades caused deficits in reaching control; here we show evidence that these effects or decrements in performance are magnified as the trial is completed reach-by-reach, decreasing reaching speed incrementally.

During visually guided reaching, eye movements are coordinated in a synchronous manner with limb movements; however, the eye acquires the target well in advance of the limb and is typically anchored to the spatial target while the reach is completed (29). This suggests that saccadic suppression may minimize potential interference for other effectors, e.g., the limb. In fact, an entire sub-field in sports medicine is devoted to “quiet” eye or limiting eye movements during ongoing limb movements. Previous studies have clearly documented saccadic programming can interfere with manual motor control depending on the feedback provided (30). A simple yet appropriate comparison between eye-hand coordination, with the eye and hand viewed as independent effector systems, may be the walking-while-talking dual task paradigm that has been studied extensively in stroke (31).

Concurrent performance of a cognitive task (talking) during a motor task (walking) typically leads to performance decrements because talking places additional demands on attentional resources, surpassing the capacity to process information used for walking. Although there are stark differences between walking-while-talking and eye-hand coordination, planning sequences of eye movements and brokering task switching in the TMT requires involves cognitive processes. Furthermore, eye movements that are decoupled from limb movements, as one might expect is occurring here sequentially given the dramatic increases in saccadic behavior, does have significant effects on the kinematics and trajectories of reaching (32).

